# Deficits in monitoring self-produced speech in Parkinson’s disease

**DOI:** 10.1101/823674

**Authors:** Henry Railo, Niklas Nokelainen, Saara Savolainen, Valtteri Kaasinen

## Abstract

**Objective:** Speech deficits are common in Parkinson’s disease, and behavioural findings suggest that the deficits may be due to impaired monitoring of self-produced speech. The neural mechanisms of speech deficits are not well understood. We examined a well-documented electrophysiological correlate of speech self-monitoring in patients with Parkinson’s disease and control participants.

**Methods:** We measured evoked electroencephalographic responses to self-produced and passively heard sounds (/a/ phonemes) in age-matched controls (N=18), and Parkinson’s disease patients who had minor speech impairment, but reported subjectively experiencing no speech deficits (N=17).

**Results:** During speaking, auditory evoked activity 100 ms after phonation (N1 wave) was less suppressed in Parkinson’s disease than controls when compared to the activity evoked by passively heard phonemes. This difference between the groups was driven by increased amplitudes to self-produced phonemes, and reduced amplitudes passively heard phonemes in Parkinson’s disease.

**Conclusions:** The finding indicates that auditory evoked activity is abnormally modulated during speech in Parkinson’s patients who do not subjectively notice speech impairment. This mechanism could play a role in producing speech deficits in as the disease progresses.

## Introduction

The majority of patients with Parkinson’s disease (PD) show speech impairments typically classified as hypokinetic dysarthria (Ho et al., 1998; Logemann et al. 1978; Sapir et al., 2001). The major speech symptoms are inaccurate articulation, variable speech rate, monotonic and hypophonic loudness of speech, and harsh voice quality. Yet, unlike primary motor symptoms of PD, speech deficits often do not respond to dopamine therapy (Louis et al. 2001; Skodda et al. 2010), and dysarthria is a well-known side-effect of deep-brain stimulation (Follett et al. 2010). Speech impairment was for a long time seen as a late symptom of PD, emerging as a by-product of the primary motor symptoms. More recent studies show that speech abnormalities are present early in the disease, and could even be considered a potential biomarker of PD (Hlavnika et al. 2017; Moreau and Pinto 2019). Yet, treatment of speech deficits remain difficult because its neural basis is not understood (Pinto et al. 2004a).

Speech requires the brain to continuously monitor how well produced actions match the desired outcome (Hickok 2012). The brain tracks how well produced actions (such as the loudness of spoken words) match the attempted outcome by relying on corollary discharge (CD) signals. According to current models, the CD signals are predictions about the sensory consequences of the produced actions which the brain communicates to sensory cortical areas (Guenther et al. 2006; Houde and Nagarajan 2011). During speech, left ventral premotor cortex (vPMC) communicates such feedforward prediction signals to auditory and somatosensory cortex. The sensory areas, in turn, feedback mismatch signals (prediction errors) back to the motor cortex, enabling rapid and efficient adjustment of speech. That is, if the produced speech does not match the attempted (i.e. predicted) speech, prediction errors enable the individual to rapidly modulate their speech volume. Similar predictive mechanisms may also contribute to speech perception (Cope et al., 2017; McClelland et al., 2006; Norris et al., 2016).

Patients with PD respond abnormally to perturbations in speech output when compared to neurologically healthy control participants. Depending on whether the fundamental frequency or first formant frequency is perturbed, the patients produce either an abnormally large (Chen et al., 2013; Liu, Wang, Metman, & Larson, 2012; Mollaei, Shiller, Baum, & Gracco, 2016) or small (Mollaei et al. 2013, 2016) correction to their speech. Some studies have not observed differences in compensation magnitude between PD patients and control participants (Kiran and Larson 2001; Abur et al. 2018). While perceptual and speech motor control abnormalities alone could in part explain the speech deficits, they could also be a sign of altered speech self-monitoring. Specifically, it has been suggested that PD patients’ brain detects a mismatch between produced and heard speech, but corrective response is not properly subsumed into subsequent speech commands (Abur et al. 2018). Deficits in speech self-monitoring is also suggested by the fact that PD patients are often not fully aware of their speech deficits (Kwan and Whitehill 2011), whereas they are typically well aware of their primary motor symptoms (Leritz et al., 2004). PD patients overestimate the volume of their own speech, and report experiencing speaking in loud voice, although their speech sound pressure level is actually reduced (Fox & Ramig, 1997; Ho ert al., 2008).

The aim of the present study was to examine the electrophysiological correlates of speech-monitoring in PD patients. Brain imaging studies have reported that activation in frontal areas during speech in PD differs from control participants. Some studies have reported decreased activation in primary or supplemental motor cortices in PD when compared to healthy control participants (Pinto et al. 2004b; Rektorova et al. 2007; Baumann et al. 2018). Also increased activation in premotor and supplementary motor areas have been reported (Liotti et al. 2003; Pinto et al. 2004b). These increased activations during speech tasks have been interpreted as *consequences* of speech deficits (compensatory processes), and patients in these studies had clear speech deficits. Arnold, Gehrig, Gispert, Seifried and Kell (2014) examined patients who did not experience speech deficits with the aim of examining if abnormal activations could reflect causes of speech deficits instead of their consequences. The results showed that when participants read passages out loud, inferior frontal, premotor and auditory cortices were more active in PD patients when compared to healthy controls. Moreover, Arnold et al. (2014) observed weaker functional connectivity between premotor cortex and auditory cortex in PD than controls during reading, suggesting weakened CD signals in PD.

Because the study by Arnold et al. (2014) did not include a passive listening control condition, it is difficult to draw conclusions about auditory speech self-monitoring. Another drawback of brain imaging studies is that they give no insight into the time-course of activity in different brain areas. Here we examined the time-course auditory evoked activity when participant either produced phonemes or listened to a recording of the same phonemes. A large body of studies shows that during speech the amplitude of auditory evoked cortical activity is suppressed when compared to a condition where the same sounds are passively listened (Curio et al. 2000; Houde et al. 2002; Ford and Mathalon 2005; Heinks-Maldonado et al. 2005; Behroozmand et al. 2011; Chang et al. 2013; Ford et al. 2013). This phenomenon is called the speaking-induced suppression (SIS), and it is assumed to reflect the influence of the CD signals on auditory cortex. In event-related potentials (ERP), SIS is strongest over central sites around 100 ms after speech/sound onset (i.e. N1 wave). Large SIS amplitudes correspond to large prediction errors: When speech output is artificially perturbed, the SIS reduces because the N1 wave in the active speech condition is increased in size (Heinks-Maldonado et al. 2005; Behroozmand and Larson 2011).

The only study that has directly examined the neural correlates of speech self-monitoring in PD is the study by Huang et al. (2016). The authors asked PD patients and control participants to sustain a vowel for 5–6 seconds, during which auditory feedback was unexpectedly modulated. The results showed that PD did not affect the amplitude of N1 (i.e. SIS), but modulated the later P2 evoked response. This finding does not indicate that PD patients do not have altered SIS, because when auditory feedback is perturbed in the middle of sustained phonation, no SIS is observed, but studies instead report an amplified P2 wave (Behroozmand et al. 2009, 2011; Chang et al. 2013). The amplified P2 is assumed to reflect a different process than SIS: whereas SIS represents suppression of initial auditory activation that matches the predictions, the amplification of P2 reflects corrections to vocal output (Hawco and Jones 2009; Scheerer and Jones 2018).

Our hypotheses were as follows. If the monitoring process where forward model predictions about speech output are compared to actual speech output is altered in PD patients, this should be reflected in the SIS amplitude. The amplitude of the SIS reflects how well the produced speech matches the predicted speech. Amplified SIS in PD compared to healthy controls would suggest that PD patients’ speech control system suppresses auditory evoked activity more than controls. Reduced SIS (in PD compared to controls), on the other hand, would indicate that the PD patients’ auditory cortex detects a higher-than-normal mismatch between produced and predicted speech. We tested patients and control participants who did not subjectively report experiencing any speech impairments.

## Methods

The experiment was approved by the Ethics committee of the hospital district of South-West Finland. A written informed consent was obtained prior the study. The study was conducted according to the principles of the Declaration of Helsinki. Participants received a small fee for participating.

### Participants

We tested 20 PD patients and 20 age-matched control participants without neurological disease. PD patients had been diagnosed using the UK Brain Bank criteria or Movement Disorder Society (MDS) clinical criteria. One control participant was left-handed, others were right-handed. The two groups were similar with respect to other measures than motor and speech deficits, which were assessed using the MDS UPDRS (Unified Parkinson’s Disease Rating Scale, part III, motor symptoms) scale. None of the patients or control participants reported experiencing any problems with speech.

Mini Mental-State (MMSE) questionnaire was used to assess cognitive impairment (Folstein et al. 1975). One PD patient was excluded from the data analysis because his MMSE-score suggested possible mild cognitive impairment (MMSE score was 23). Beck’s Depression Inventory was used to asses depression (Beck 1976). One PD patient was excluded due to relatively high score (27 total score) in BDI. Finally, one patient had to be excluded because of a technical failure during EEG recording, and two control participants were excluded because of massive artifacts in EEG. Therefore, the final sample was 17 patients and 18 control participants. Comparison of the demographical information, and clinical and neuropsychological measurements are presented in Table 1.

**Table 1.**
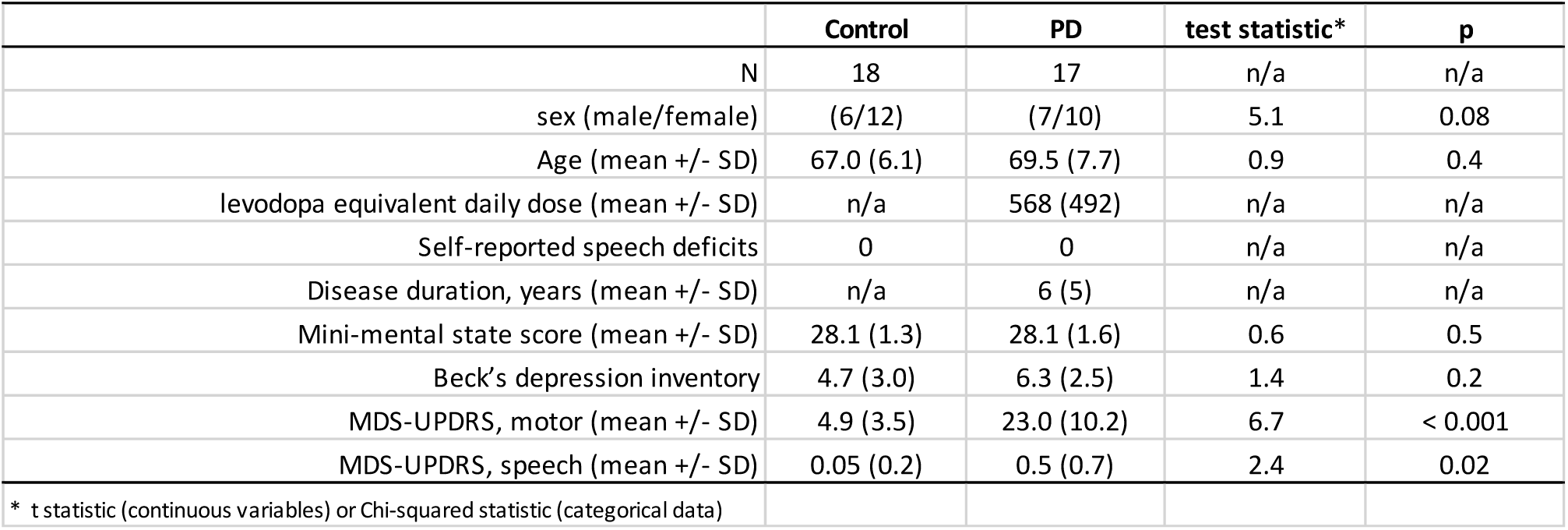
Comparison of demographic, clinical and neuropsychological data between PD and control groups.

|  |  |  |  |  |
| --- | --- | --- | --- | --- |
|  | Control | PD | test statistic* | p |
| N | 18 | 17 | n/a | n/a |
| sex (male/female) | (6/12) | (7/10) | 5.1 | 0.08 |
| Age (mean +/- SD) | 67.0 (6.1) | 69.5 (7.7) | 0.9 | 0.4 |
| levodopa equivalent daily dose (mean +/- SD) | n/a | 568 (492) | n/a | n/a |
| Self-reported speech deficits | 0 | 0 | n/a | n/a |
| Disease duration, years (mean +/- SD) | n/a | 6 (5) | n/a | n/a |
| Mini-mental state score (mean +/- SD) | 28.1 (1.3) | 28.1 (1.6) | 0.6 | 0.5 |
| Beck's depression inventory | 4.7 (3.0) | 6.3 (2.5) | 1.4 | 0.2 |
| MDS-UPDRS, motor (mean +/- SD) | 4.9 (3.5) | 23.0 (10.2) | 6.7 | < 0.001 |
| MDS-UPDRS, speech (mean +/- SD) | 0.05 (0.2) | 0.5 (0.7) | 2.4 | 0.02 |
| * t statistic (continuous variables) or Chi-squared statistic (categorical data) |  |  |  |  |

Eleven patients were on levodopa medication, and sixteen were treated with MAO-inhibitors and/or dopamine agonists. One patient was on anti-cholinergic medication. The mean levodopa equivalent dose across the patients was 568 mg (SD = 492 mg). The patients were asked to go through a voluntary medication break before the test session to counteract possible intervening effects of medication. Twelve patients underwent the medication break, and did not take their morning medication during the day of the study (at least 12 h break in medication). These twelve patients can therefore be considered being in OFF phase (the remaining patients are in ON phase, and did not go through the medication brake).

### Task and procedure

In the psychophysical task the participants listened to /a/ phonemes from a loudspeaker (Creative T40 Series II), and they were asked to repeat the phoneme with a matching volume. There were five different volume levels, ranging from weak to loud speaking volumes (on average 67 dB), but otherwise the phonemes were identical. This procedure was repeated 65 times in total (13 times for each volume level). The participant’s response was measured using a microphone (Vivitar TVM-1) positioned halfway between the loudspeaker and the participant. Response amplitude was determined by calculating the mean absolute amplitude of samples that were higher than 50% of the maximum absolute amplitude. Before each test session the position of the microphone was calibrated to ensure that volume was measured accurately. This calibration consisted of playing an identical phoneme as a response from a loudspeaker positioned where the participants head should be. This procedure yielded near perfect correspondence between stimulus and response, verifying that the volume measurement was accurate. No EEG was measured during this task. Our hypothesis was that if the patients show hypophonic speech they should systematically produce lower volume responses than control participants, as shown by Clark et al. (2014).

In the EEG condition, the participant was first asked to utter the phoneme /a/ approximately every 5 seconds for 10 minutes with their normal speaking volume while minimizing facial movements (“Speaking condition”). Vocalizations lasted on average approximately 350 ms, and patients/participants produced on average 120 (SD = 61) vocalizations during the 10 min recording. The participants wore headphones with a microphone. White noise and the participant’s own speech was played back through the headphones. In the second condition (“Listening condition”) the same phonemes they had just spoken were played back to them through the headphones (again with white noise in the background). With the white noise added to suppress direct perceptions of one’s own speech, the volume of the spoken/listened phonemes could be matched. In both Speak and Listening conditions the white noise was identical, and played throughout the recording (i.e. not just during vocalization).

To accurately measure the onset time of the phonemes, the signal from the loudspeakers of the headphones was recorded by the EEG amplifier. The auditory signal is thus measured by the same system as brain activity, which means that the two signals are perfectly synchronized. Markers were inserted to the EEG data to the time-points that corresponded to the onsets of phonemes following a semi-automated procedure during off-line analysis. First, only the signal from the headphones was selected and high-pass filtered at 100 Hz; because only high-frequency information is present, the transient onset of sound could be easily detected. Second, if the signal remained above a predefined threshold for minimum of 10 samples, a marker was added to the timepoint where the threshold was first crossed. All markers and trials were inspected by the researcher, and any trials where the onset of the sound was unclear, were excluded from the analysis.

### Electroencephalography

EEG was measured with 64 active electrodes with NeurOne Tesla amplifier. Sampling rate was 500 Hz. In addition, EOG was measured from electrodes placed beside and below the right eye. EEG was processed using EEGLAB v14.1.1 software (Delorme and Makeig 2004) on Matlab 2014b. Channels were first automatically inspected, and bad channels interpolated. The channel used to measure the auditory signal was removed before EEG analysis. EEG was fist high-pass (0.1 Hz) and then low-pass (40 Hz) filtered. Speech related and other EEG artefacts were removed using artifact subspace reconstruction (ASR). ASR resembles principle component analysis (PCA) in that data is reconstructed after first excluding artifactual components. ASR rejects components automatically by determining thresholds based on clean segments of EEG data. Here, ASR was utilized instead of PCA or independent component analysis (ICA) because large speech-related artefacts remained in the data after PCA or ICA-based artifact rejection. For technical details of ASR, the reader is referred to (Mullen et al. 2015). After ASR, data was epoched, removed electrodes were interpolated, and the data referenced to linked mastoids. Epochs containing abnormal data were rejected using pop_jointprob function. This left on average 105 (SD = 53) trials per condition per participant for the EEG analyses.

### Statistical analysis

The results of the psychophysical task were analyzed using linear mixed-effects model. The model included the intensity of the test stimulus, group, and their interaction as predictors. Control group was coded as 0 (i.e. baseline), and the PD group as 1.

ERPs were analyzed using Condition x Group linear regression models. In the model the control group and Listening condition were the reference categories. The statistical analysis was performed in a mass univariate approach: ERPs of all channels and time points between 0 and 400 ms were analyzed. To reduce the number of statistical tests, the analysis was performed in 10 ms steps (the average of 5 consecutive EEG samples was analyzed). We used a non-parametric permutation approach with 1000 repetitions to test for statistical significance (Maris and Oostenveld 2007). To take into account the fact that statistically significant effects cluster in time and space, threshold free cluster enhancement (TFCE) was employed (Smith and Nichols 2009). TFCE eliminates the need for researcher-defined clustering thresholds. TFCE works by modulating the inputted statistical map (here a map of t values) based on how clustered the data is. When a given pixel is part of a cluster of pixels with similar effect, TFCE amplifies the values. TFCE was performed on the real dataset and permuted data where the observations were randomly shuffled. To control for multiple comparisons, null distribution was obtained by selecting from each permutation iteration the maximum TFCE value (Maris and Oostenveld 2007). TFCE values whose p value < .01 were considered statistically significant effects. TFCE was performed using the Limo EEG package (Pernet et al. 2011).

The datasets are available for download at https://osf.io/eys97/.

## Results

### Speech deficits, and the psychophysical task

While most participants and PD patients were assessed to not have speech deficits according to the item representing speech deficit in MDS-UPDRS, PD patients were observed to have speech deficits more frequently than control participants (Table 1). Six of the 17 patients were rated with some speech deficits in the UPDRS (i.e. score 1 or 2).

Our hypothesis was that even though patients reported no speech deficits, weak hypophonia would be detectable using psychophysical assessment. Both, patients and control participants produced higher volume speech to louder test stimuli, as shown in Figure 1. In general, for low volume phonemes participants/patients tended to produce a higher volume sound than the test stimulus, whereas for higher sound volumes they tended to underestimate the loudness. When analyzed using Test Stimulus Amplitude × Group linear mixed-effects regression (df = 1696) where individual participants’ intercepts are allowed to vary randomly, the main effect of Test Stimulus Amplitude strongly predicted their response (β = 0.49, CI = [0.45–0.52], t = 28.05, p < .0001), and the interaction between Test Stimulus Amplitude and Group also reached statistical significance (β = - 0.08, CI = [−0.13–-0.035], t = −3.36, p = 0.0008). This interaction indicates that, on average, the influence of Test Stimulus Amplitude (i.e. slope) was smaller in PD patients. However, as seen in Figure 1, this effect is influenced by few individual patients, and when the between-subject variation in the slope is taken into account as a random-effect factor (in addition to intercept), the interaction is not statistically significant (β = −0.068, CI = [−0.20–0.06], t = −0.99, p = 0.32).

**Figure 1.**
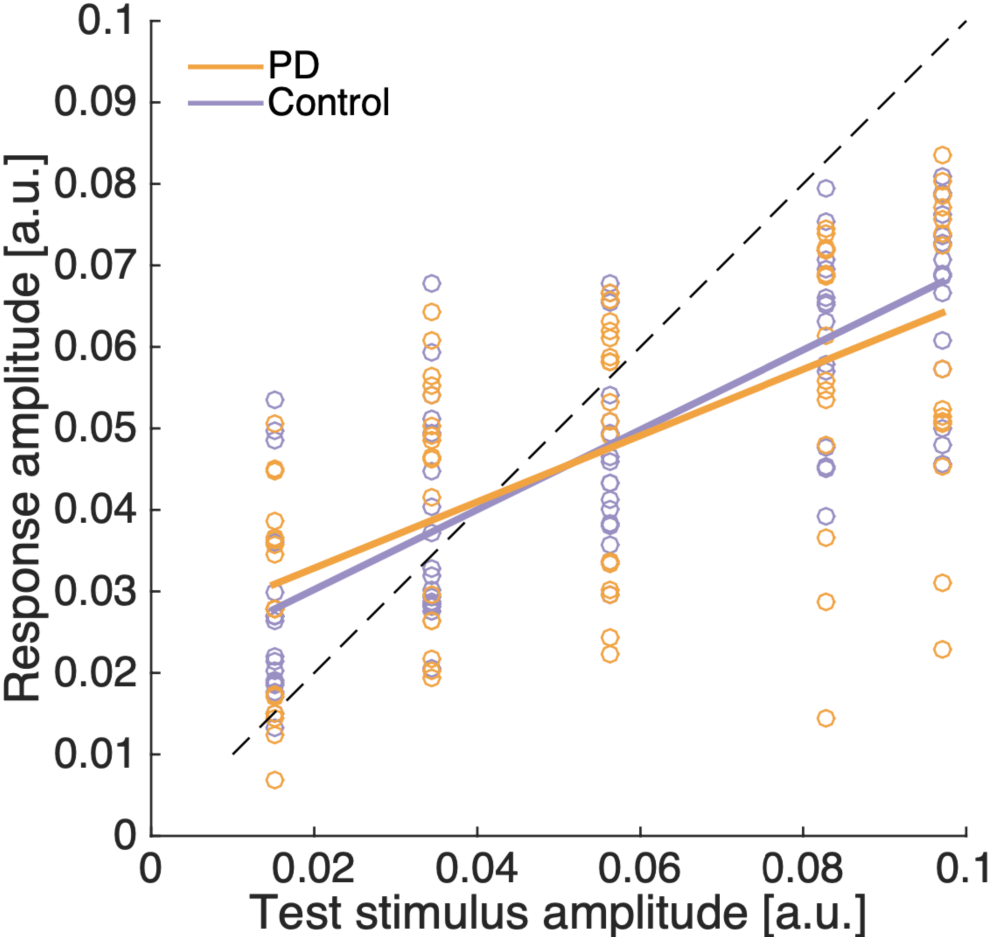
Results of the psychophysical task. PD patients and control participants are depicted as different colors (PD = orange). Dots represent individual participants/patients. The solid lines show the fixed-effects results of the linear model (Test Stimulus Amplitude × Group linear mixed-effects model, with by-subject intercepts). The dashed line is a reference line that denotes perfect correspondence between test stimulus and response amplitude.

### EEG

As shown in Figure 2A-C, ERPs time-locked to the onset of the spoken (Speaking condition; Fig. 2B) or heard (Listening condition, Fig. 2C) phoneme were calculated. In both conditions, N1 (negative peak at 80 ms), and P2 (positive peak at 190 ms) peaks are observed. The topography of the ERPs in the Speak and Listening condition is shown in Figure 2D-E (averaged across patients and control participants).

**Figure 2.**
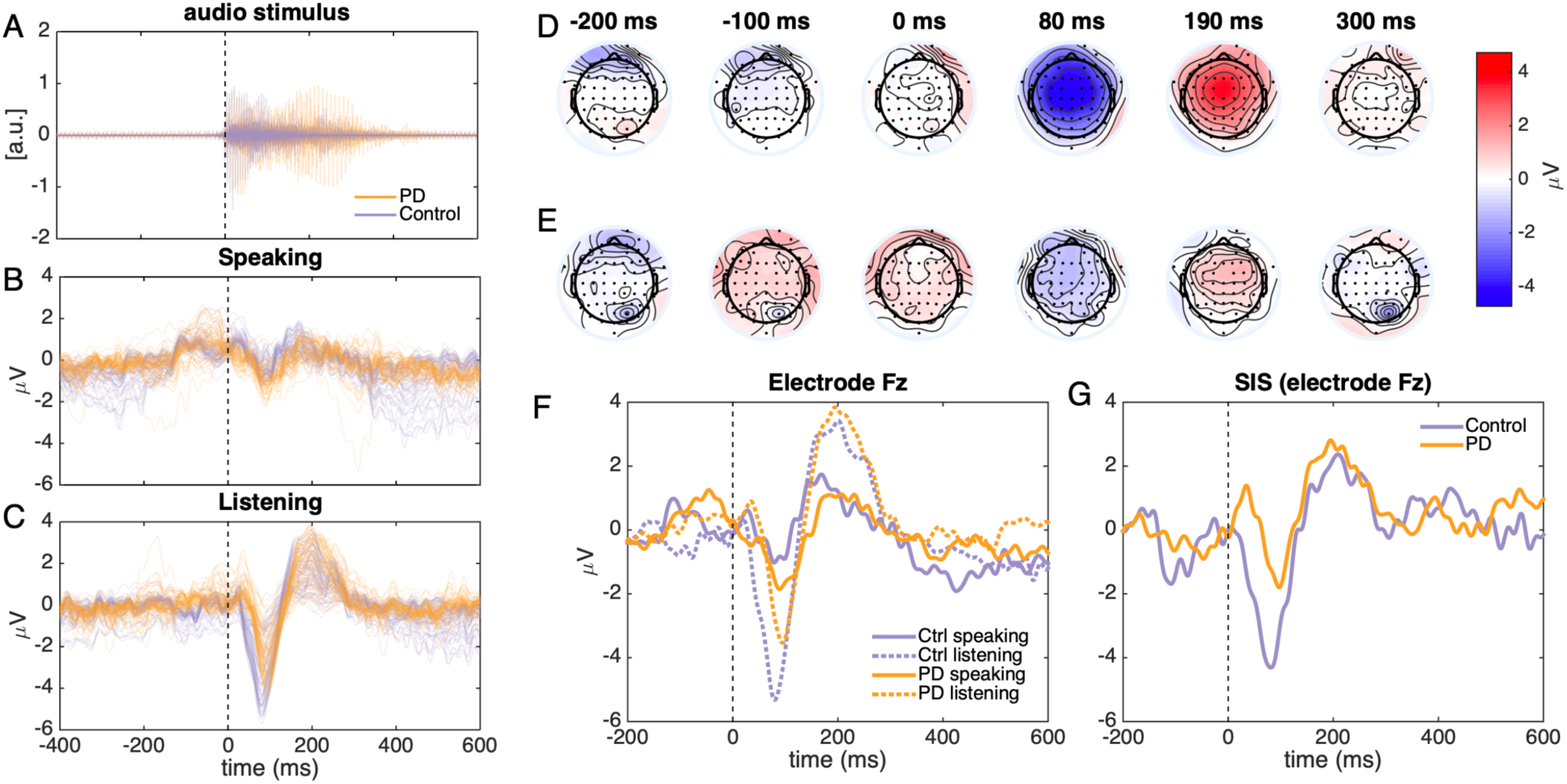
Auditory ERPs. A) The auditory signal recorded by EEG (average over all participants/patients). B) A butterfly plot of ERPs in the Speaking condition. C) Plot of ERPs in the Listening condition. Each line depicts one channel. In each panel the orange line is PD patients, and blue line is the control group. Panels D and E show the scalp distribution of ERP in the listening and speaking conditions, respectively (averaged over the two groups). F) Shows the ERPs from electrode Fz, and G) the corresponding speaking-induced suppression (SIS: Listening minus Speaking condition) in the control and PD groups.

The results of the statistical analysis (Condition x Group mass univariate regression) are presented in Figure 3. The panels show the contribution of each predictor on the ERP amplitude at each time point and channel (color denotes t value). The leftmost panel shows the intercept of the regression model, which corresponds to Listening condition in the control group. The blue and red areas represent the N1 and P2 ERP waves, respectively. The second panel from left shows how ERPs in the Listening condition in the control group change during vocalization. Positive t values in the N1 and negative t values in the P2 time-window indicate that the amplitude of these waves are reduced in the Speaking condition relative to Listening condition. This effect is the speaking induced suppression (SIS), and it is assumed to reflect the influence of CD signals on auditory processing: the motor cortex communicates top-down predictions (CD) about the upcoming speech to the auditory cortex, and hence evoked neural activity is weaker. As previously, this effect was strongest in central electrode locations. ERPs and corresponding SIS from channel Fz are shown in Fig. 2F and Fig. 2G.

**Figure 3.**
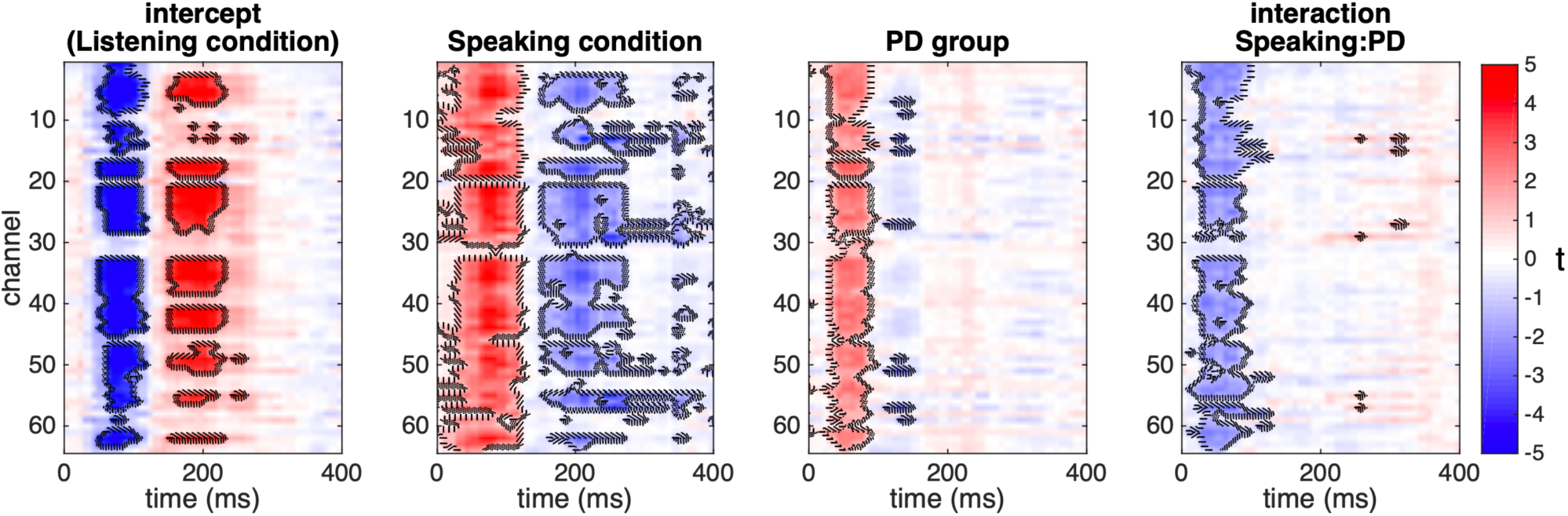
Results of the mass univariate regression models. Channels are presented on y axis and time on x axis. Colors denote t values. Strong colors that are surrounded by a contour are statistically significant (p < .001) clusters of effects as determined by permutation testing and TFCE.

The panel entitled “PD group” in Fig. 3 shows how ERPs in the Listen condition change when the participant belongs to the PD group: N1 amplitude are reduced, as demonstrated by positive t values. This effect is also clearly seen in Figure 2F. Finally, the rightmost panel of Fig. 3 shows the interaction between condition and group: Negative t values in the N1 time-window show that, when compared to the control group, the decrease in N1 amplitude during Speaking condition is smaller in the PD group. In sum, in the N1 time-window, PD patients showed both reduced amplitudes to passively heard phonemes, and reduced suppression of actively spoken phonemes, when compared to control participants. The reduction in SIS in PD patients is evident when it is expressed in proportion of N1 amplitude in the Listening condition: In the control group the peak amplitude of the N1 wave at electrode Fz was reduced by 81% during vocalization whereas in the PD group the suppression was only 49% of peak N1 amplitude.

As shown in Fig. 2G, the PD and control groups show a pronounced difference in SIS at very early latency. In fact, whereas control participants show a statistically significant SIS already 35 ms after phonation (t = −2.1, p = .045), the PD patients show an opposite effect (speaking induced *amplification*) at this point (t = 2.4, p = .028). Fig. 2C suggests that this effect could be caused by a delayed N1 wave in the PD group in the Listening condition. To examine this, we statistically analyzed N1 peak latencies using linear regression. As shown in Fig. 4A, N1 peak latency in the Listening condition strongly predicted peak SIS latency (β = .74, t = 3.7, p = .0008); this effect was not statistically significantly different between the groups. The PD and control groups did not statistically significantly differ with respect to N1 peak latency in the Listening condition (p = .71), N1 latency in the Speaking condition (p =. 13), or with respect to SIS peak latency (p = .71). Altogether this suggests while SIS peak latency is strongly predicted by the peak latency of the N1 in the Listening condition, the difference in SIS amplitude is cannot be attributed to a delayed N1 peak in PD (instead, difference between PD and control groups in SIS is driven by decreased amplitude of the Listening N1 wave in PD).

**Figure 4.**
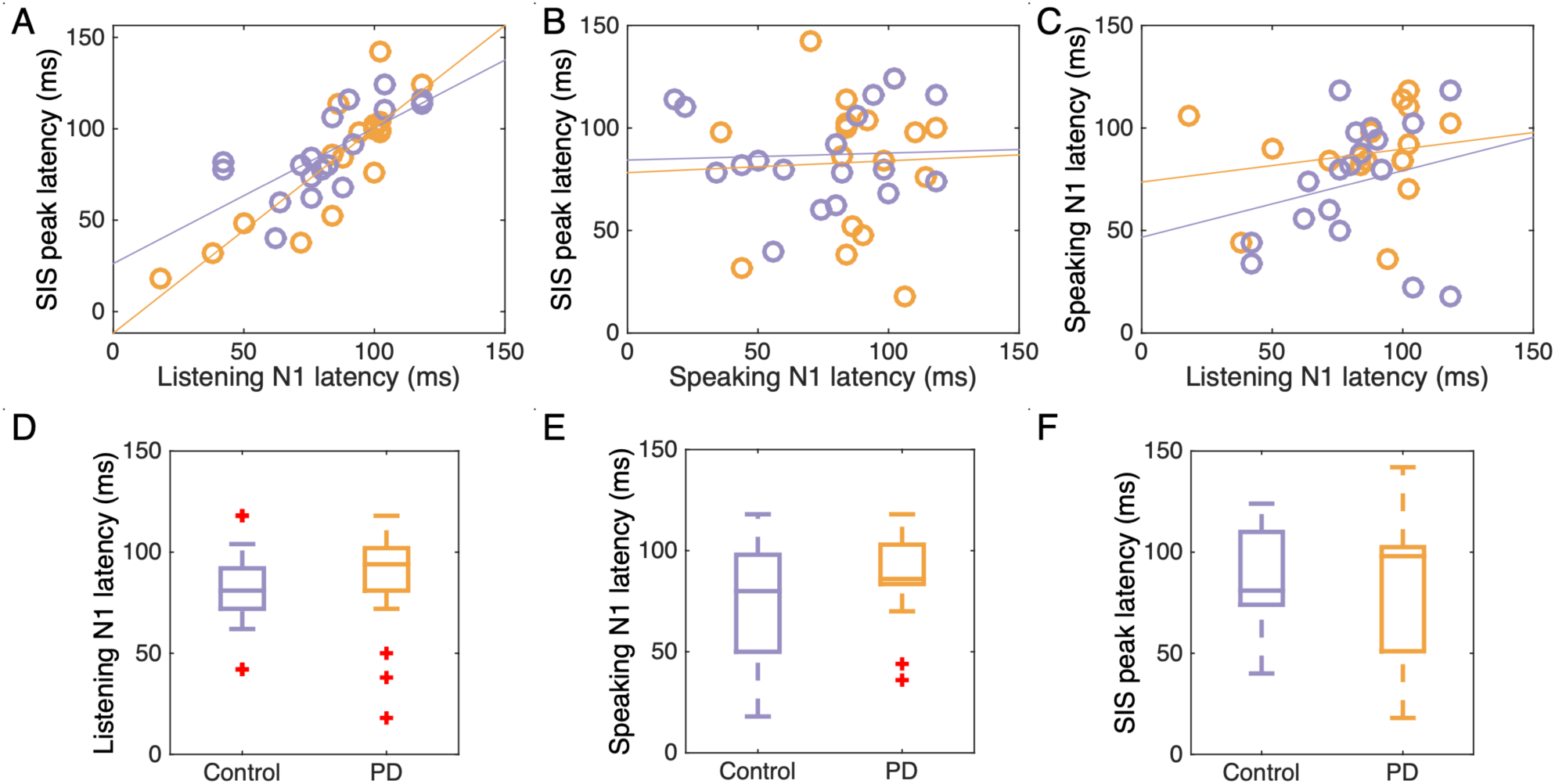
N1 peak latency analysis results. A) SIS peak latency is strongly predicted by N1 peak latency in the Listening condition. B) In contrast, N1 peak latency in the Speaking condition did not predict SIS peak latency. C) N1 peak latency in the Listening and Speaking condition did not correlate. The PD patients and control participants did not show statistically significant differences in D) N1 peak latency in Listening condition, E) N1 peak latency in Speaking condition, or F) SIS.

## Discussion

Models of speech production (Guenther et al. 2006; Houde and Nagarajan 2011) assume that self-produced sounds are compared to predicted speech output during early cortical processing stages. Mismatch between predicted and produced speech are sent from the auditory cortex to motor cortex. A well-known electrophysiological correlate of this process is the SIS (Curio et al. 2000; Houde et al. 2002; Heinks-Maldonado et al. 2005; Behroozmand et al. 2011; Chang et al. 2013; Ford et al. 2013). The results of the present study show reduced SIS in PD patients relative to control group, suggesting that in PD patients auditory cortex detected a higher than normal mismatch between predicted and produced speech. The patients themselves reported experiencing no speech impairments, although neurological examination (UPDRS speech item) revealed a weak speech impairment.

The difference between the groups in SIS amplitude was observed within the first 120 ms after phonation (i.e. N1 wave). Whereas the SIS onset in the control participants is on average around 20 ms after phoneme onset, in PD patients SIS onset is clearly delayed (average onset around 75 ms). Because of this delay, auditory evoked activity during speaking is actually amplified, not suppressed, right after phonation in the PD group relative to the listening condition. This difference between the groups was not related to variation in N1 peak latency.

The reduced SIS is PD patients when compared to control participants was influenced by two factors. First, the PD patients showed reduced N1 amplitudes to passively heard phonemes, and second, amplitudes to spoken phonemes were less suppressed in PD patients relative to control participants. That N1 amplitude in the Listening condition was reduced in PD suggests that deficits in monitoring self-produced speech could be influenced by deficits auditory processing. While most previous auditory ERP studies report that later evoked potentials are modulated by PD (Tanaka et al. 2000; Solís-Vivanco et al. 2015; Huang et al. 2016; Cavanagh et al. 2018), PD patients have been reported to be less sensitive to detect first formant differences during passive listening of vowels (Mollaei et al. 2019).

Although N1 amplitude in the Listening condition was suppressed in PD patients, N1 amplitudes in the Speaking condition were less suppressed when compared to control participants. According to one interpretation, the suppression of auditory cortical activity by CD signals allows individuals to distinguish self-produced sounds from external sounds. Based on this, our findings suggest that PD patients have deficits in tracking self-produced speech. Alternatively, the reduced SIS could be interpreted as a mismatch (prediction error) between intended and produced speech output (Chang et al. 2013). In normal speech, this type of feedback from auditory cortex is assumed to fine-tune motor speech control. In PD the reduced SIS could lead to a situation where speech motor control becomes dominated by the feedback from auditory cortex, which could then contribute to speech deficits.

Huang et al. (2016) observed that PD and control groups did not differ with respect to SIS/N1 amplitudes, but PD patients made stronger corrections to perturbed speech, which was reflected in enhanced P2 amplitudes. Our results are not at odds with the findings by Huang et al. (2016) but extend them. The present results show that when a paradigm that is sensitive to SIS is used, PD patients show reduced SIS amplitudes in the N1 time-window when compared to control participants. Our observation that PD patients have reduced SIS (i.e. increased mismatch signaling) are consistent with the observation that PD patients tend to produce stronger corrections to speech when fundamental frequency is perturbed (Liu et al. 2012; Chen et al. 2013; Huang et al. 2016; Mollaei et al. 2016).

Because our patients did not show strong speech deficits, the link between altered SIS and speech deficits in PD remains to be demonstrated. Instead of reflecting a specific correlate of speech deficit, the reduced SIS could be a sign of a more general deficit in neural processing. Railo et al. (2018) showed that in a double-saccade task used to probe self-monitoring in the oculomotor system PD patients made errors whose direction suggested that the patients overestimated the amplitude of their saccades. Furthermore, reduced SIS has also been reported in psychotic patients when compared to healthy controls (Ford and Mathalon 2005; Perez et al. 2012; Ford et al. 2013).

Because PD medication was not systematically manipulated in the present study, potential confounds produced by the medication cannot be ruled out. However, the strength of the present study is that the patients showed very weak speech deficits and cognitive decline, which makes the comparison to control participants more straightforward. That is, it is unlikely that the observed changes in ERPs would simply consequences of speech deficits.

## Conclusion

We have provided evidence that the electrophysiological correlate of speech self-monitoring (SIS) is altered in PD. Consistent with previous reports, the present finding suggests that abnormalities in speech-related neural activity occur early in PD in patients who do not display or experience strong speech deficits (e.g. Moreau & Pinto, 2019). Our results indicate that PD patients’ auditory system detects a higher-than-normal mismatch (corresponding to reduced SIS) between planned and produced speech. This mechanism could contribute to speech deficits as the disease progresses.

## Authors’ roles

H.R. designed the experiment, supervised data collection, analyzed the data, and wrote the first manuscript draft. N.N and S.S. collected the data. V.K. helped to organize the research, supervised data collection, and reviewed and revised the manuscript.

## Acknowledgements

The study (H.R.) was funded by Academy of Finland (grant #308533).

## References

Abur D, Lester-Smith RA, Daliri A, Lupiani AA, Guenther FH, Stepp CE. Sensorimotor adaptation of voice fundamental frequency in Parkinson’s disease. PLoS One. 2018; https://doi.org/10.1371/journal.pone.0191839

Arnold C, Gehrig J, Gispert S, Seifried C, Kell CA. Pathomechanisms and compensatory efforts related to Parkinsonian speech. NeuroImage Clin. 2014; 4: 82–97.

Baumann A, Nebel A, Granert O, Giehl K, Wolff S, Schmidt W, et al. Neural Correlates of Hypokinetic Dysarthria and Mechanisms of Effective Voice Treatment in Parkinson Disease. Neurorehabil Neural Repair. 2018; 32(12):1055–1066.

Beck AT. Cognitive therapy and the emotional disorders. Intunivpress, New York. 1976.

Behroozmand R, Karvelis L, Liu H, Larson CR. Vocalization-induced enhancement of the auditory cortex responsiveness during voice F0 feedback perturbation. Clin Neurophysiol. 2009; 120(7):1303–12. doi: 10.1016/j.clinph.2009.04.022

Behroozmand R, Larson CR. Error-dependent modulation of speech-induced auditory suppression for pitch-shifted voice feedback. BMC Neurosci. 2011; 6;12:54. doi: 10.1186/1471-2202-12-54.

Behroozmand R, Liu H, Larson CR. Time-dependent neural processing of auditory feedback during voice pitch error detection. J Cogn Neurosci. 2011; 23(5):1205–17. doi: 10.1162/jocn.2010.21447.

Cavanagh JF, Kumar P, Mueller AA, Richardson SP, Mueen A. Diminished EEG habituation to novel events effectively classifies Parkinson’s patients. Clin Neurophysiol. 2018; 129(2):409–418. doi: 10.1016/j.clinph.2017.11.023.

Chang EF, Niziolek CA, Knight RT, Nagarajan SS, Houde JF. Human cortical sensorimotor network underlying feedback control of vocal pitch. Proc Natl Acad Sci U S A. 2013; 110(7):2653–8. doi: 10.1073/pnas.1216827110.

Chen X, Zhu X, Wang EQ, Chen L, Li W, Chen Z, et al. Sensorimotor control of vocal pitch production in Parkinson’s disease. Brain Res. 2013;

Clark JP, Adams SG, Dykstra AD, Moodie S, Jog M. Loudness perception and speech intensity control in Parkinson’s disease. J Commun Disord. 2014; 1646:269–277. doi: 10.1016/j.brainres.2016.06.013.

Cope TE, Sohoglu E, Sedley W, Patterson K, Jones PS, Wiggins J, et al. Evidence for causal top-down frontal contributions to predictive processes in speech perception. Nat Commun. 2017; 8(1):2154. doi: 10.1038/s41467-017-01958-7.

Curio G, Neuloh G, Numminen J, Jousmäki V, Hari R. Speaking modifies voice-evoked activity in the human auditory cortex. Hum Brain Mapp. 2000; 9(4):183–91.

Delorme A, Makeig S. EEGLAB: An open source toolbox for analysis of single-trial EEG dynamics including independent component analysis. J Neurosci Methods. 2004; 134(1):9–21.

Follett KA, Weaver FM, Stern M, Hur K, Harris CL, Luo P, et al. Pallidal versus subthalamic deep-brain stimulation for Parkinson’s disease. N Engl J Med. 2010; 362(22):2077–91. doi: 10.1056/NEJMoa0907083.

Folstein MF, Folstein SE, McHugh PR. “Mini-mental state”. A practical method for grading the cognitive state of patients for the clinician. J Psychiatr Res. 1975; 12(3):189–98.

Ford JM, Mathalon DH. Corollary discharge dysfunction in schizophrenia: Can it explain auditory hallucinations? Int J Psychophys. 2005; 58(2-3):179–89.

Ford JM, Mathalon DH, Roach BJ, Keedy SK, Reilly JL, Gershon ES, et al. Neurophysiological evidence of corollary discharge function during vocalization in psychotic patients and their nonpsychotic first-degree relatives. Schizophr Bull. 2013; 39(6):1272–80. doi: 10.1093/schbul/sbs129.

Fox CM, Ramig LO. Vocal Sound Pressure Level and Self-Perception of Speech and Voice in Men and Women with Idiopathic Parkinson Disease. Am J Speech-Language Pathol. 1997; 6, 85–94. https://doi.org/10.1044/1058-0360.0602.85

Guenther FH, Ghosh SS, Tourville JA. Neural modeling and imaging of the cortical interactions underlying syllable production. Brain Lang. 2006; 96(3):280–301.

Hawco CS, Jones JA. Control of vocalization at utterance onset and mid-utterance: Different mechanisms for different goals. Brain Res. 2009; 1276:131–9. doi: 10.1016/j.brainres.2009.04.033.

Heinks-Maldonado TH, Mathalon DH, Gray M, Ford JM. Fine-tuning of auditory cortex during speech production. Psychophysiology. 2005; 42(2):180–90.

Hickok G. Computational neuroanatomy of speech production. Nature Reviews Neuroscience. 2012. 13(2):135–45. doi: 10.1038/nrn3158.

Hlavnika J, Cmejla R, Tykalová T, Šonka K, Ruzicka E, Rusz J. Automated analysis of connected speech reveals early biomarkers of Parkinson’s disease in patients with rapid eye movement sleep behaviour disorder. Sci Rep. 2017; 7(1):12. doi: 10.1038/s41598-017-00047-5.

Ho AK, Bradshaw JL, Iansek R. For better or worse: The effect of Levodopa on speech in Parkinson’s disease. Mov Disord. 2008; 23(4):574–80. doi: 10.1002/mds.21899.

Ho AK, Iansek R, Marigliani C, Bradshaw JL, Gates S. Speech impairment in a large sample of patients with Parkinson’s disease. Behav Neurol. 1998; 11(3):131–137.

Houde JF, Nagarajan SS. Speech production as state feedback control. Frontiers in Human Neuroscience. 2011. 5:82. doi: 10.3389/fnhum.2011.00082. eCollection 2011.

Houde JF, Nagarajan SS, Sekihara K, Merzenich MM. Modulation of the auditory cortex during speech: An MEG study. J Cogn Neurosci. 2002; 14(8):1125–38.

Huang X, Chen X, Yan N, Jones JA, Wang EQ, Chen L, et al. The impact of parkinson’s disease on the cortical mechanisms that support auditory–motor integration for voice control. Hum Brain Mapp. 2016; 4248–4261. doi: 10.1002/hbm.23306.

Kiran S, Larson CR. Effect of duration of pitch-shifted feedback on vocal responses in patients with Parkinson’s disease. J Speech, Lang Hear Res. 2001; 44(5):975–87.

Kwan LC, Whitehill TL. Perception of speech by individuals with Parkinson’s disease: A review. Parkinson’s Disease. 2011. 389767. doi: 10.4061/2011/389767.

Leritz E, Loftis C, Crucian G, Friedman W, Bowers D. Self-awareness of deficits in Parkinson disease. Clin Neuropsychol. 2004; 18(3):352–61.

Liotti M, Ramig LO, Vogel D, New P, Cook CI. Hypophonia in Parkinson’s disease Neural correlates of voice treatment revealed by PET. Neurology. 2003; 60(3):432–40.

Liu H, Wang EQ, Metman LV, Larson CR. Vocal responses to perturbations in voice auditory feedback in individuals with parkinson’s disease. PLoS One. 2012; e33629. doi: 10.1371/journal.pone.0033629.

Logemann JA, Fisher HB, Boshes B, Blonsky ER. Frequency and cooccurrence of vocal tract dysfunctions in the speech of a large sample of Parkinson patients. J Speech Hear Disord. 1978; 43(1):47–57.

Louis ED, Winfield L, Fahn S, Ford B. Speech dysfluency exacerbated by levodopa in Parkinson’s disease. Mov Disord. 2001; 16(3):562–5.

Maris E, Oostenveld R. Nonparametric statistical testing of EEG- and MEG-data. J Neurosci Methods. 2007; 164(1):177–90.

McClelland JL, Mirman D, Holt LL. Are there interactive processes in speech perception? Trends Cogn Sci. 2006; 10(8):363–9.

Mollaei F, Shiller DM, Baum SR, Gracco VL. Sensorimotor control of vocal pitch and formant frequencies in Parkinson’s disease. Brain Res. 2016; 1646:269–277. doi: 10.1016/j.brainres.2016.06.013.

Mollaei F, Shiller DM, Baum SR, Gracco VL. The relationship between speech perceptual discrimination and speech production in parkinson’s disease. J Speech, Lang Hear Res. 2019; 62(12):4256–4268. doi: 10.1044/2019_JSLHR-S-18-0425.

Mollaei F, Shiller DM, Gracco VL. Sensorimotor adaptation of speech in Parkinson’s disease. Mov Disord. 2013; 28(12):1668–74. doi: 10.1002/mds.25588.

Moreau C, Pinto S. Misconceptions about speech impairment in Parkinson’s disease. Mov Disord. 2019;34(10):1471–5. 34(10):1471-1475. doi: 10.1002/mds.27791.

Mullen TR, Kothe CAE, Chi YM, Ojeda A, Kerth T, Makeig S, et al. Real-time neuroimaging and cognitive monitoring using wearable dry EEG. IEEE Trans Biomed Eng. 2015; 62(11):2553–67. doi: 10.1109/TBME.2015.2481482.

Norris D, McQueen JM, Cutler A. Prediction, Bayesian inference and feedback in speech recognition. Lang Cogn Neurosci. 2016; 31(1):4–18.

Perez VB, Ford JM, Roach BJ, Loewy RL, Stuart BK, Vinogradov S, et al. Auditory cortex responsiveness during talking and listening: Early illness schizophrenia and patients at clinical high-risk for psychosis. Schizophr Bull. 2012; 38(6):1216–24. doi: 10.1093/schbul/sbr124.

Pernet CR, Chauveau N, Gaspar C, Rousselet GA. LIMO EEG: A toolbox for hierarchical linear modeling of electroencephalographic data. Comput Intell Neurosci. 2011; 2011:831409. doi: 10.1155/2011/831409.

Pinto S, Ozsancak C, Tripoliti E, Thobois S, Limousin-Dowsey P, Auzou P. Treatments for dysarthria in Parkinson’s disease. Lancet Neurology. 2004a. 3(9):547–56.

Pinto S, Thobois S, Costes N, Le Bars D, Benabid AL, Broussolle E, et al. Subthalamic nucleus stimulation and dysarthria in Parkinson’s disease: A PET study. Brain. 2004b; 127(Pt 3):602–15.

Railo H, Olkoniemi H, Eeronheimo E, Pääkkönen O, Joutsa J, Kaasinen V. Dopamine and eye movement control in Parkinson’s disease: Deficits in corollary discharge signals? PeerJ. 2018; 6:e6038. doi: 10.7717/peerj.6038.

Rektorova I, Barrett J, Mikl M, Rektor I, Paus T. Functional abnormalities in the primary orofacial sensorimotor cortex during speech in Parkinson’s disease. Mov Disord. 2007; 22(14):2043–51.

Sapir S, Pawlas AA, Ramig LO, Countryman S, O’Brien C, Hoehn MM, et al. Voice and Speech Abnormalities in Parkinson Disease: Relation to Severity of Motor Impairment, Duration of Disease, Medication, Depression, Gender, and Age. J Med Speech Lang Pathol. 2001; 9(4), 231–226.

Scheerer NE, Jones JA. The role of auditory feedback at vocalization onset and mid-utterance. Front Psychol. 2018; 9:2019. doi: 10.3389/fpsyg.2018.02019.

Skodda S, Visser W, Schlegel U. Short- and long-term dopaminergic effects on dysarthria in early Parkinson’s disease. J Neural Transm. 2010; 117(2):197–205. doi: 10.1007/s00702-009-0351-5.

Smith SM, Nichols TE. Threshold-free cluster enhancement: Addressing problems of smoothing, threshold dependence and localisation in cluster inference. Neuroimage. 2009; 44(1):83–98. doi: 10.1016/j.neuroimage.2008.03.061.

Solís-Vivanco R, Rodríguez-Violante M, Rodríguez-Agudelo Y, Schilmann A, Rodríguez-Ortiz U, Ricardo-Garcell J. The P3a wave: A reliable neurophysiological measure of Parkinson’s disease duration and severity. Clin Neurophysiol. 2015; 126(11):2142–9. doi: 10.1016/j.clinph.2014.12.024.

Tanaka H, Koenig T, Pascual-Marqui RD, Hirata K, Kochi K, Lehmann D. Event-related potential and EEG Parkinson’s disease without and with dementia. Dement Geriatr Cogn Disord. 2000; 11(1):39–45.

